# Cortisol associated with hypometabolism across the Alzheimer’s disease spectrum

**DOI:** 10.1101/514968

**Authors:** Miranka Wirth, Catharina Lange, Willem Huijbers, for the Alzheimer’s Disease Neuroimaging Initiative

**Affiliations:** Charité - Universitätsmedizin Berlin, Corporate Member of Freie Universität Berlin, Humboldt-Universität zu Berlin, and Berlin Institute of Health, Klinik und Hochschulambulanz für Neurologie, Berlin, Germany; Charité - Universitätsmedizin Berlin, Corporate Member of Freie Universität Berlin, Humboldt-Universität zu Berlin, and Berlin Institute of Health, NeuroCure Cluster of Excellence, Berlin, Germany; Charité - Universitätsmedizin Berlin, corporate member of Freie Universität Berlin, Humboldt-Universität zu Berlin, and Berlin Institute of Health, Center for Stroke Research, Berlin, Germany; Charité - Universitätsmedizin Berlin, Corporate Member of Freie Universität Berlin, Humboldt-Universität zu Berlin, and Berlin Institute of Health, Department of Nuclear Medicine, Berlin, Germany; Tilburg University, Department of Cognitive Science and Artificial Intelligence, Jheronimus Academy of Data Science, Tilburg, the Netherlands

**Keywords:** stress, risk factor, hypothalamic-pituitary-adrenal axis, neurodegeneration, ADNI

## Abstract

**Objective:** Hypothalamic-pituitary-adrenal (HPA) dysregulation is proposed as a risk factor for Alzheimer’s disease (AD). This cross-sectional study assessed relationships between plasma cortisol levels and neuroimaging biomarkers, specifically brain glucose metabolism and gray matter volume, across the AD spectrum.

**Methods:** Cognitively normal older adults and patients with mild cognitive impairment (MCI) and AD dementia were included from the Alzheimer’s Disease Neuroimaging Initiative. Participants (n = 556) were selected based on availability of baseline measures of plasma cortisol levels and gray matter volume, as estimated with magnetic resonance imaging. Within a subsample (n = 288), we examined brain glucose metabolism (n = 288) as with positron emission tomography. Relationships between plasma cortisol and AD neuroimaging biomarkers were assessed using regions-of-interest and voxel-wise analyses.

**Results:** Across the entire cohort, higher plasma cortisol was also related to lower gray matter volume, most notably in the left lateral temporal-parietal-occipital regions. Importantly, higher plasma cortisol concentration was also related to hypometabolism, especially in lateral temporo-parietal and medial parietal regions. When stratified by diagnosis, these negative associations were most pronounced in MCI and AD patients.

**Interpretation:** High plasma cortisol was associated with hypometabolism predominantly in AD-sensitive regions. Our results indicate that HPA axis activation could influence brain metabolism and exacerbate existing AD pathological processes. This is consistent with a notion that stress is a conceivable target for intervention to slow down AD progression. Future studies should delineate underlying pathological mechanisms and investigate if clinical or lifestyle interventions could alleviate negative actions of stress on AD.

## 1 Introduction

Stress, mediated by the hypothalamic-pituitary-adrenal (HPA) axis, has been proposed as a risk factor for the development of Alzheimer’s disease (AD) ^1^. Known as the most important human glucocorticoid released by HPA axis activation, cortisol modulates homeostatic body functions, including negative feedback mechanisms in the brain ^2^. Elevated levels of circulating cortisol over time may, however, prompt brain injury and increase vulnerability to neurodegenerative diseases ^3-5^.

There is evidence that HPA axis activation and dysregulation may aggravate pathological processes related to AD. Cortisol levels, commonly measured in blood plasma/serum, salvia, urine or cerebrospinal fluid (CSF), are increased in patients with AD and amnestic MCI ^6, 7^. Moreover, higher cortisol levels were previously related to gray matter atrophy in the hippocampus ^8-11^ across the AD spectrum, from cognitively normal older adults, to mild cognitive impairment (MCI) and clinical AD. These brain changes may contribute to cognitive deficits ^9, 10, 12^ and higher risk of clinical progression ^7, 13, 14^ associated with stress hormone elevation.

By contrast, effects of circulating cortisol on cerebral glucose metabolism are largely unknown in the course of AD pathogenesis. Decreased glucose metabolism within frontal-temporal-parietal association cortices, measured using positron emission tomography (PET), is a key feature of prodromal and clinical AD ^15-17^. Chronic exposure to increased glucocorticoid concentration was formerly linked to impaired glucose metabolism in the hippocampus and other brain areas in rat models ^18^ as well as in hypercortisolemic patients with Cushing syndrome ^19, 20^. Given these observations, we hypothesize that brain glucose metabolism may be affected by the actions of stress hormones along the AD spectrum.

The present cross-sectional study aimed to determine the relationship between plasma cortisol levels and cerebral glucose metabolism, measured using 2-[F-18]-fluoro-2-deoxy-D-glucose (FDG) Positron Emission Tomography (PET), across the AD spectrum. First, we assessed the effect of plasma cortisol on gray matter volume, measured using structural magnetic resonance imaging (MRI), with the aim to replicate previous findings ^8^. Secondly, we investigated the unexplored relationship between circulating cortisol and FDG-PET. The methodological approach was chosen to highlight specific regional patterns of cortisol-associated effects in each imaging modality.

## 2 Methods

Cross-sectional data used in this study were obtained from the Alzheimer’s Disease Neuroimaging Initiative (ADNI) database (adni.loni.usc.edu). The ADNI was launched in 2003 as a public-private partnership, led by Principal Investigator Michael W. Weiner, MD. The primary goal of ADNI has been to test whether serial magnetic resonance imaging (MRI), positron emission tomography (PET), other biological markers, and clinical and neuropsychological assessment can be combined to measure the progression of mild cognitive impairment (MCI) and early Alzheimer’s disease (AD). For up-to-date information, see www.adni-info.org.

### 2.1 Participants

Participants were selected from the ADNI database (ADNI I cohort) with regard to availability of following assessments: (i) baseline plasma cortisol concentration, (ii) baseline structural (high-resolution T1-weighted) MRI, and baseline cerebral FDG PET (subgroup). Neuroimaging data was restricted to image data acquired within a time window of ±100 days anchored to the clinical baseline assessment (incl. collection of plasma cortisol data). The selection flowchart is depicted in Figure 1.

**Figure 1.**
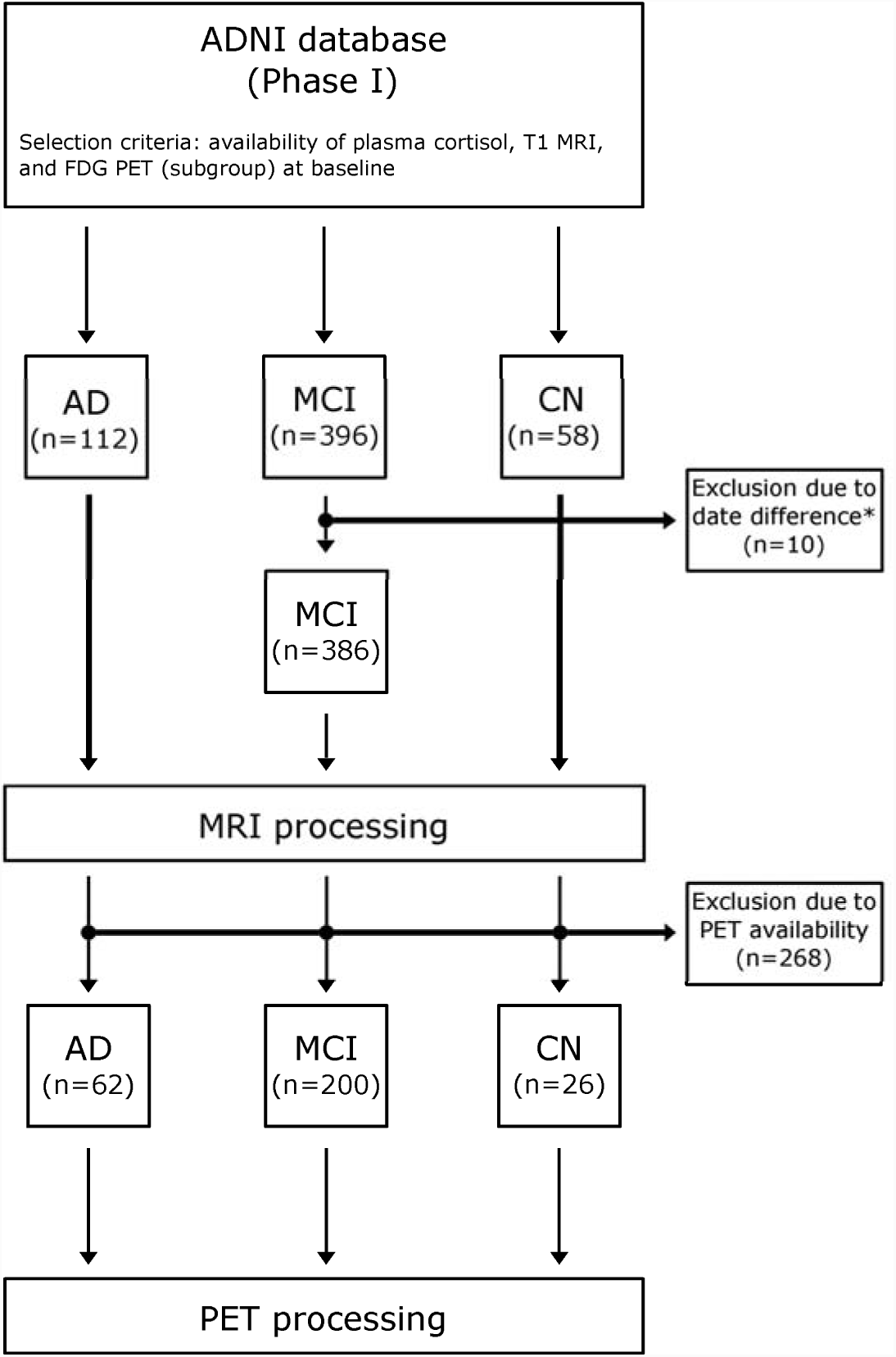
Participant selection flowchart. The graphs depict the selection procedure from the ADNI database. *Selection of neuroimaging data was restricted to images acquired within a time window of ±100 days anchored to clinical baseline examinations. Abbreviations – AD: Alzheimer’s disease dementia, ADNI: Alzheimer’s Disease Neuroimaging Initiative, FDG: 2-[F-18]-fluoro-2-deoxy-D-glucose, MCI: mild cognitive impairment, MRI: magnetic resonance imaging, CN: participants with normal cognition.

The final sample included in this study had 556 participants with diagnostic labels of early dementia due to AD (n = 112), MCI in later stages (n = 386), and normal cognition (CN: n = 58). A subsample of 288 participants had FDG PET available (AD: n = 62, MCI: n = 200, and CN: n = 26).

Details on inclusion/exclusion criteria are available on the ADNI website (http://adni.loni.usc.edu/methods/). Briefly, at the time of study enrollment, participants are between 55 to 90 (inclusive) year old. The CN participants have mini-mental state examination (MMSE) ^21^ scores between 24 to 30 (inclusive), a Clinical Dementia Rating (CDR) Scale ^22^ of 0, no signs of depression, no memory complaints, and normal memory function. Participant with MCI have MMSE score between 24 to 30 (inclusive), a CDR of 0.5, subjective memory complaints, objective memory dysfunction, and preserved general cognition and functional performance. Participants with AD demonstrate initial MMSE scores between 20 to 26 (inclusive), a CDR of 0.5 or 1.0, and meet the NINCDS-ADRDA criteria for clinically probable AD ^23^.

Standard protocol approvals, registrations, and patient consents. Ethics approval was obtained by the ADNI investigators. All study participants provided written informed consent.

### 2.2 Acquisition of data

#### 2.2.1 Plasma sample collection and analysis

Plasma cortisol data was provided by the Biomarkers Consortium Plasma Proteomics Project and extracted using the ADNImerge package. Essays and quantification methods are described in detail in the data primer, available at (http://adni.loni.usc.edu/methods/) and explained in previous publications ^8, 24^. In brief, baseline plasma cortisol was quantified in overnight fasting blood samples collected from each participant. Whole-blood samples were collected into 10 ml BD lavender top K2EDTA coated vacutainers, centrifuged within one hour after collection. Immediately after, blood plasma was transferred to a polypropylene transfer tube and placed in dry ice. Samples were shipped to the ADNI Biomarker Core Laboratory at the University of Pennsylvania. Following thawing at room temperature, 0.5 mL aliquots were prepared from plasma samples and stored in polypropylene aliquot tubes at −80 C until analyzed. Plasma samples were interrogated by Rules-Based Medicine, Inc. (RBM, Austin, TX) in accordance with ADNI standard operating procedures.

#### 2.2.2 Acquisition of MRI images

MRI acquisition protocols were set-up to harmonize image quality across MR hardware platforms ^25^. In particular, high-resolution T1-weighted MR images were acquired using a sagittal 3-dimensional magnetization prepared rapid gradient echo (3D-MPRAGE) sequence with an approximate TR = 2400 ms, minimum full TE, TI = 1000 ms, and flip angle of 8° (scan parameters vary between sites, scanner platforms, and software versions).

#### 2.2.3 Acquisition of FDG PET images

Cerebral FDG PET data was acquired on multiple PET scanners of varying resolution, but platform-specific acquisition and reconstruction protocols were used to harmonize image quality across all ADNI centers ^26^. FDG PET was performed according to a dynamic protocol resulting in 6 frames, each with a 5 min duration from 30 to 60 min post injection.

### 2.3 Preprocessing of structural MRI images

In general, image processing and voxel-wise statistical testing was conducted in Matlab environment (version R2013b, The Mathworks, Natick, USA). All images were downloaded from the ADNI repository as “unpreprocessed” data. Mcverter was used for DICOM-to-Nifti-conversion (https://lcni.uoregon.edu/downloads/mriconvert/mriconvert-and-mcverter).

For brain tissue segmentation into gray matter (GM), white matter (WM), and CSF, the unified segmentation algorithm (Statistical Parametric Mapping, Wellcome Trust Centre for Neuroimaging, London, UK, www.fil.ion.ucl.ac.uk/spm, version SPM12) was deployed with default parameters, except that image data was resampled to 2×2×2 mm ^27,28^. Normalization to the Montreal Neurological Institute (MNI) template space was performed using diffeomorphic anatomical registration through exponentiated lie algebra (DARTEL) with default parameters and registration to ‘existing templates’ ^29^. The IXI555 templates, which are defined in MNI space and provided by the CAT12 toolbox, were used (http://dbm.neuro.uni-jena.de/vbm). Atlas-based hippocampal volumetry as well as voxel-wise statistical testing was performed on normalized and modulated gray matter images processed by DARTEL.

Hippocampal volume was calculated using the previously published masks according the EADC-ADNI Harmonized Protocol for hippocampal segmentation (HarP; probabilistic labels thresholded at 100% overlap) and inclusion of both, modulated gray matter and white matter voxel values ^30^. Total intracranial volume (TIV) was estimated by the method of Keihaninejad and colleagues ^31^. Hippocampal volumes were corrected for TIV using a simple ratio method.

In order to prepare voxel-wise multiple regression analyses, the gray matter volume images were smoothed by a three-dimensional (3D) Gaussian kernel with full width at half maximum (FWHM) of 6 mm.

### 2.4 Preprocessing of FDG PET images

Reconstructed dynamic (or static, if dynamic was not available) PET data was downloaded from the ADNI repository in its original image format (“as archived”, DICOM, Interfile, or ECAT). Image data was converted to Nifti, i.e. from DICOM and ECAT using SPM12 or from Interfile using ImageConverter (version 1.1.5, Turku PET center).

Preprocessing was performed fully automatically using a custom-made pipeline described elsewhere ^32^, except that the PET data was spatially normalized to MNI space using the parameter estimates from the MRI (see section 2.3). The relative standardized uptake value (SUVr) was calculated voxel-wise using the whole brain parenchyma as reference region.

Mean FDG SUVr was evaluated using a regions-of-interest (ROI) template, comprising neocortical areas that known to be hypometabolic in AD. Specifically, the FDG-based AD template included parietal, temporal, and frontal regions and was defined previously from voxel-wise comparisons of FDG PET images in an largely independent ADNI sample of CN and AD patients (except for five overlapping individuals) ^32^.

In order to prepare voxel-wise multiple regression analyses, SUVr images were smoothed by a 3D Gaussian kernel with FWHM of 12 mm.

### 2.5 Statistics

In general, statistical analyses of sample characteristics and ROI analysis were performed with IBM SPSS Statistics (version 22, IBM Corp., Armonk, NY, USA). Whole brain data was visualized using Python (3.6.0) and the nibabel (DOI: 10.5281/zenodo.1464282) and nilearn (version 0.5.0b) modules. Scatterplots were generated using R (version 3.5.1) and ggplot2 ^33^.

#### 2.5.1 Sample characteristics

The entire sample and each diagnostic groups were characterized using baseline demographic, clinical, and biomarker characteristics, as extracted from the ADNImerge data (R package) downloaded on April 17^th^ 2018 ^34^.

Following clinical and neuropsychological measures were selected for sample description: individual baseline scores from the CDR Sum of Boxes (CDR-SB), the MMSE, the Rey auditory verbal learning test (RAVLT), percent forgetting subscale ^35^, and the Geriatric Depression Scale (GDS) ^36^. The entire sample and diagnostic groups were further characterized using the Apolipoprotein E number of ε4 alleles (APOE4 status), the average FDG SUVr within AD-sensitive regions, total hippocampal volume, and baseline plasma cortisol levels.

One-way analysis of variance (ANOVA) models were computed with diagnostic group as independent factor and the respective sample characteristics as dependent variable. In case of a significant effect between groups, post-hoc multiple comparisons were performed using either the Tukey’s honestly significant difference test (assuming equal variance) or the Games Howell test (unequal variance).

#### 2.5.2 Regions-of interest analysis

A regions-of interest (ROI) analysis was performed in pre-defined ROIs representing regions that are sensitively affected by hypometabolism and gray matter atrophy in AD patients. For structural MRI, we assessed bilateral hippocampal volume according HarP ^30^, corrected for TIV using a simple ratio method. For FDG, we examined the mean SUVr in the combined AD ROI described above ^32^.

Pearson correlation analysis was performed to evaluate relationships between plasma cortisol and each AD-sensitive ROI for the entire sample. Follow-up stratification by diagnostic group was conducted to explore respective relationships within each diagnostic group. To evaluate possible group differences, one-way ANOVA models were computed with each AD-sensitive ROI as dependent variable and diagnostic group (CN, MCI, and AD), plasma cortisol level, and their interactive term as independent factors. Scatter plots were created to visualize the relationships between plasma cortisol and glucose metabolism as well as gray matter volume measured in each pre-selected AD ROI, respectively.

#### 2.5.3 Voxel-wise analysis

Voxel-wise regression analysis was run under SPM12 to evaluate relationships between plasma cortisol and imaging modalities (FDG PET and structural MRI) in the entire sample. Results were evaluated for significance at a False Discovery Rate (FDR) adjusted p value of < 0.05 and a minimum cluster size of 20. Voxel-wise analyses were adjusted for age and TIV (note: gray matter volume only) and restricted to cerebral gray matter using an explicit mask (cerebellum excluded).

For follow-up analysis, average FDG SUVr and total gray matter volume were delineated from combined clusters of significant negative associations between plasma cortisol and the respective imaging modality (here referred as cortisol meta ROI). For MRI, the cortisol meta ROI values were corrected for TIV using a simple ratio method. Pearson correlation analyses were performed to estimate associations between plasma cortisol and each cortisol meta ROI for the respective diagnostic groups. To explore possible group differences, one-way ANOVA models were computed with each cortisol meta ROI as dependent variable and diagnostic group (CN, MCI, and AD), plasma cortisol, and their interactive term as independent factors.

Scatter plots were created to visualize the relationships between plasma cortisol and glucose metabolism as well as gray matter volume measured in each cortisol meta ROI, respectively.

#### 2.5.4 Adjustment procedure

Second-level statistical analyses within each cortisol meta ROI respective both imaging modalities, FDG and MRI, were carried out with adjustments for appropriate covariates as follows: Using correlation analyses, we assessed preliminary relationships between age, education, gender, subclinical depression (measured using the GDS), APOE status, and respective ROIs (dependent variable). In case of a considerable correlation (*p* < 0.1) of any covariate with the dependent variable of interest, partial correlation analyses were conducted including the particular covariate(s).

## 3 Results

### 3.1 Characteristics of participants

Descriptive demographic, clinical, and biomarker characteristics are provided for the entire sample and within each diagnostic group in Table 1. Diagnostic groups were comparable in age, gender and education. As expected, CN, MCI, and AD groups differed significantly in clinical measures, the APOE4 status, and the selected AD-sensitive ROIs for FDG SUVr and gray matter volume. Plasma cortisol levels were significantly increased in AD patients compared to the MCI group.

**Table 1.** Baseline demographics, clinical, and biomarker characteristics.

| Characteristics | Entire cohort | CN | MCI | AD | p-value<sup>†</sup> |
| --- | --- | --- | --- | --- | --- |
| n [total] | 556 | 58 | 386 | 112 |  |
| **Demographics** |  |  |  |  |  |
| age [years] | 74.8 (7.4) | 75.1 (5.8) | 74.7 (7.4) | 74.8 (8.0) | 0.942 |
| gender [no. females] | 213 | 28 | 138 | 47 | 0.126 |
| education [years] | 15.6 (3.0) | 15.7 (2.8) | 15.7 (3.0) | 15.1 (3.2) | 0.185 |
| **Clinical measures** |  |  |  |  |  |
| CDR-SB | 2.0 (1.6) | 0.0 (0.1) | 1.6 (0.9) | 4.3 (1.6) | <0.001<sup>a,b,c</sup> |
| MMSE | 26.6 (2.4) | 28.9 (1.2) | 27.1 (1.8) | 23.6 (1.9) | <0.001<sup>a,b,c</sup> |
| RAVLT (% forgetting) | 68.1 (33.3), n = 553 | 34.6 (33.9) | 67.8 (31.4), n = 385 | 86.5 (24.2), n = 110 | <0.001<sup>a,b,c</sup> |
| GDS | 1.5 (1.4) | 0.9 (1.2) | 1.6 (1.4) | 1.7 (1.34) | <0.001<sup>b,c</sup> |
| **Biomarkers** |  |  |  |  |  |
| APOE4 status 0 / 1 / 2 [n: ε4 alleles] | 271 / 216 / 69 | 53 / 5 / 0 | 182 / 158 / 46 | 36 / 53 / 23 | <0.001<sup>a,b,c</sup> |
| MRI hippocampus [ml] | 5.18 (0.89) | 5.84 (0.59) | 5.23 (0.84) | 4.68 (0.93) | <0.001<sup>a,b,c</sup> |
| FDG AD template | 1.01 (0.04), **n = 288** | 1.04 (0.02), **n = 26** | 1.02 (0.03), **n = 200** | 0.98 (0.05), **n = 62** | <0.001<sup>a,b,c</sup> |
| Plasma cortisol [ng/ml] | 2.2 (0.1) | 2.17 (0.14) | 2.16 (0.13) | 2.21 (0.11) | **0.002<sup>a</sup>** |

Numbers are given if applicable as mean and standard deviation (parenthesis). The actual sample size is given if different from sample size specified in first row.
**Abbreviations** – AD: participants with dementia due to Alzheimer's disease, ApoE: Apolipoprotein E, CDR-SB: clinical dementia rating (sum of boxes), CN: participants with normal cognition, FDG: 2-[F-18]-fluoro-2-deoxy-D-glucose, n: number; GDS: geriatric depression scale, MCI: participants with mild cognitive impairment, MMSE: mini-mental state examination, MRI: magnetic resonance imaging, RAVLT: Rey auditory verbal learning test.
<sup>†</sup> Statistical comparisons of numerical characteristics across diagnostic groups was performed by one-way ANOVA. In case of a significant effect between diagnostic groups, post-hoc multiple comparisons were performed using either the Tukey's honestly significant difference test (assuming equal variance) or the Games Howell test (unequal variance). Chi-square statistic was used to compare the distribution of categorical variables between the three diagnostic groups. In case of a significant effect between groups, a chi-square test was applied to all possible pairs of groups. A significance level of $p \leq 0.05$ was assumed for between all groups tests as well as for post hoc tests.
<sup>a</sup> Post-hoc analysis: significant group difference between AD and MCI.
<sup>b</sup> Post-hoc analysis: significant group difference between AD and CN.
<sup>c</sup> Post-hoc analysis: significant group difference between MCI and CN.

### 3.2 Results of ROI analyses

#### 3.2.1 MRI ROI

Higher plasma cortisol was significantly related to lower hippocampal volume in the entire sample (Figure 2 and Table 2A, correlation analysis). When stratified by diagnostic group, the significant negative relationship between plasma cortisol and hippocampal volume was present only in the MCI group. There was no significant interaction between diagnostic group and plasma cortisol in the hippocampal ROI (*F* [2,550] = 1.446, *p* = 0.236, n = 556).

**Table 2.** Relationship between plasma cortisol, gray matter volume, and glucose metabolism.

|  | Entire cohort |  |  | CN |  |  | MCI |  |  | AD |  |  | p-value<sup>†</sup> |
| --- | --- | --- | --- | --- | --- | --- | --- | --- | --- | --- | --- | --- | --- |
|  | *n* | *rho* | *p-value* | *n* | *rho* | *p-value* | *n* | *rho* | *p-value* | *n* | *rho* | *p-value* |  |
| **A. Pre-selected ROI** |  |  |  |  |  |  |  |  |  |  |  |  |  |
| Correlation analysis |  |  |  |  |  |  |  |  |  |  |  |  |  |
| MRI hippocampus (% TIV) | 556 | **-0.143** | **0.001\*\*\*** | 58 | -0.058 | 0.666 | 386 | **-0.170** | **0.001\*\*\*** | 112 | 0.012 | 0.899 | 0.236 |
| FDG AD template | 288 | **-0.199** | **0.001\*\*\*** | 26 | **0.421** | **0.032\*** | 200 | **-0.158** | **0.025\*** | 62 | **-0.256** | **0.045\*** | **0.025\*** |
| Partial correlation analysis<sup>††</sup> |  |  |  |  |  |  |  |  |  |  |  |  |  |
| MRI hippocampus (% TIV) | 556 | **-0.100** | **0.018\*** | 58 | -0.005 | 0.974 | 386 | **-0.123** | **0.016\*** | 112 | -0.026 | 0.793 | 0.548 |
| FDG AD template | 288 | **-0.199** | **0.001\*\*\*** | 26 | 0.347 | 0.105 | 200 | **-0.143** | **0.045\*** | 62 | **-0.295** | **0.023\*** | **0.021\*** |
| **B. Cortisol meta ROI** |  |  |  |  |  |  |  |  |  |  |  |  |  |
| Correlation analysis |  |  |  |  |  |  |  |  |  |  |  |  |  |
| MRI cortisol meta ROI (% TIV) | 556 | **-0.301** | **0.001\*\*\*** | 58 | **-0.434** | **0.001\*\*\*** | 386 | **-0.289** | **0.001\*\*\*** | 112 | **-0.214** | **0.024\*** | 0.747 |
| FDG cortisol meta ROI | 288 | **-0.274** | **0.001\*\*\*** | 26 | -0.180 | 0.379 | 200 | **-0.256** | **0.001\*\*\*** | 62 | -0.223 | 0.082<sup>‡</sup> | 0.596 |
| Partial correlation analysis<sup>††</sup> |  |  |  |  |  |  |  |  |  |  |  |  |  |
| MRI cortisol meta ROI (% TIV) | 556 | **-0.285** | **0.001\*\*\*** | 58 | **-0.430** | **0.001\*\*\*** | 386 | **-0.261** | **0.001\*\*\*** | 112 | **-0.215** | **0.024\*** | 0.601 |
| FDG cortisol meta ROI | 288 | **-0.277** | **0.001\*\*\*** | 26 | -0.230 | 0.279 | 200 | **-0.248** | **0.001\*\*\*** | 62 | -0.250 | 0.054<sup>‡</sup> | 0.564 |

**Abbreviations** – AD: participants with dementia due to Alzheimer's disease, CN: participants with normal cognition, FDG: 2-[F-18]-fluoro-2-deoxy-D-glucose, MCI: participants with mild cognitive impairment, MRI: magnetic resonance imaging, ROI: regions-of-interest.
<sup>†</sup> Statistical comparisons across groups was performed by one-way ANOVA with each ROI as dependent variable and diagnostic group (CN, MCI, and AD), plasma cortisol, and their interactive term as independent factors. P-values for the interactive term are provided. Significance levels: \*\*\*p ≤ 0.001, \*\*p ≤ 0.01, \*p ≤ 0.05, ‡p ≤ 0.1.
<sup>††</sup> Statistical models adjusted for covariates (see text for details).

**Figure 2.**
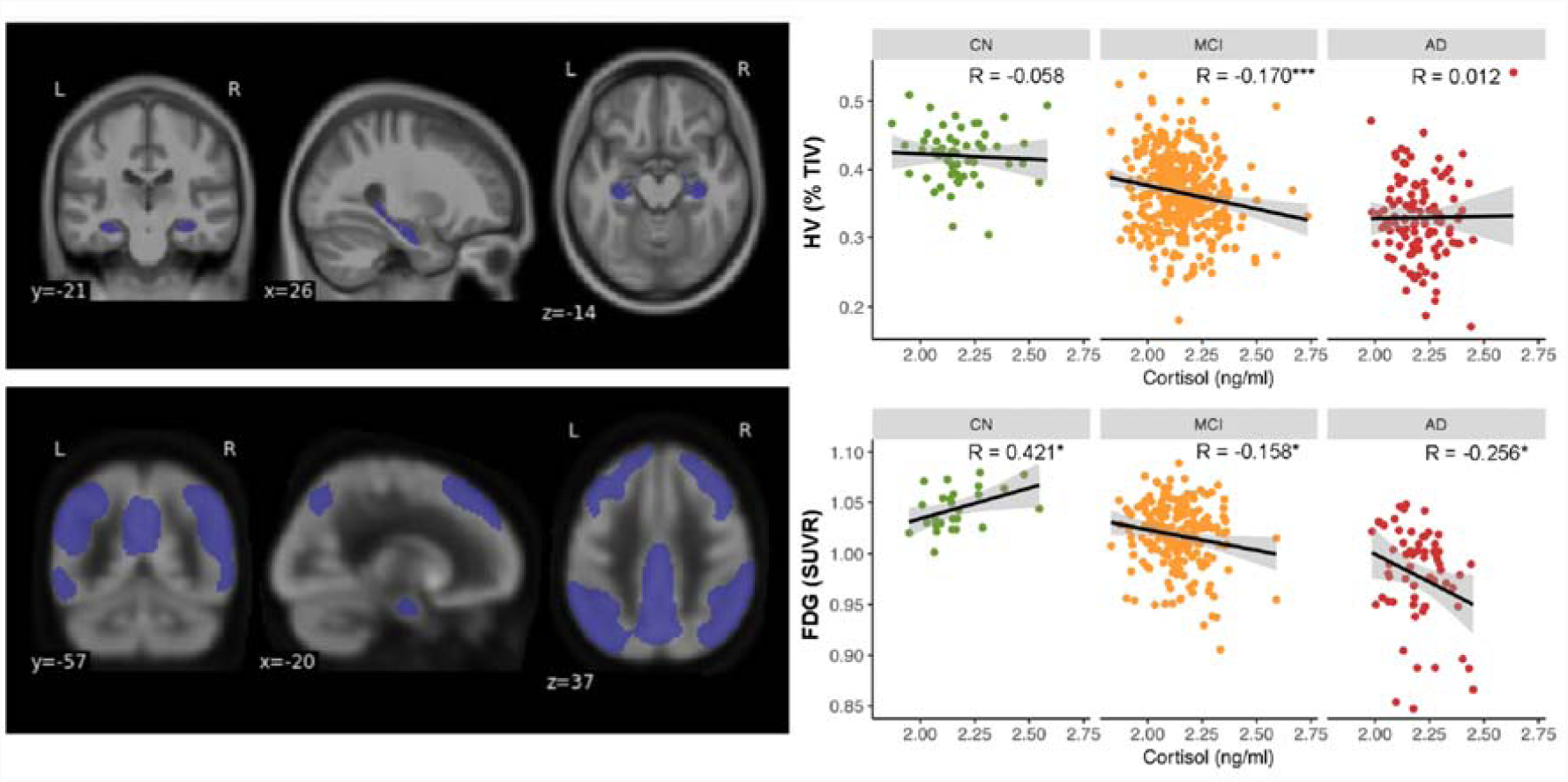
Relationship between plasma cortisol, hippocampal volume and glucose metabolism in pre-selected regions-of-interest. Top panel shows hippocampal volume (HV). Scatter plots (top right side) depict relationships between plasma cortisol and total HV (measured as % total intracranial volume [TIV]). Lower panel shows glucose metabolism (FDG) from an a-priori template, sensitive to AD pathology. Scatter plots (lower right side) depict relationships between plasma cortisol and average FDG relative standardized uptake value (SUVr). Dots represent individual data points, with cognitively normal [CN] in green, mild cognitive impairment [MCI] in orange and Alzheimer disease [AD] patients in red. Lines indicate linear trends within each diagnostic group and gray area the standard deviation. Correlation coefficients are provided in the top right for each diagnostic group. Significance levels: ***p ≤ 0.001, **p ≤ 0.01, * p ≤ 0.05, ^‡^ p ≤ 0.1.

Adjustment for covariates (age, education, gender, subclinical depression, and APOE4) did not change the results with regard to the entire sample and diagnostic groups (Table 2A, partial correlation analysis).

#### 3.2.2 FDG ROI

Higher plasma cortisol levels were significantly related to lower glucose metabolism in the FDG AD template for the entire sample (Figure 2 and Table 2A, correlation analysis). When stratified by diagnostic group, significantly negative relationships between higher plasma and glucose metabolism were seen only in the MCI and AD groups (Table 2). The interaction between diagnostic groups and plasma cortisol was significant (*F* [2,282] = 3.749, *p* = 0.025, n = 288).

Adjustment for covariates (age, subclinical depression, and APOE4) did not essentially change the results with regard to the entire sample and diagnostic groups (Table 2A, partial correlation analysis).

### 3.3 Results of voxel-wise analyses

#### 3.3.1 MRI

Higher plasma cortisol was related to lower gray matter volume, mainly in lateral temporal, temporo-parietal and occipital regions of the left hemisphere, across the entire sample (Figure 3, Table 3). There was no significant positive relationship between plasma cortisol levels and gray matter volume at the given statistical threshold.

**Table 3.**
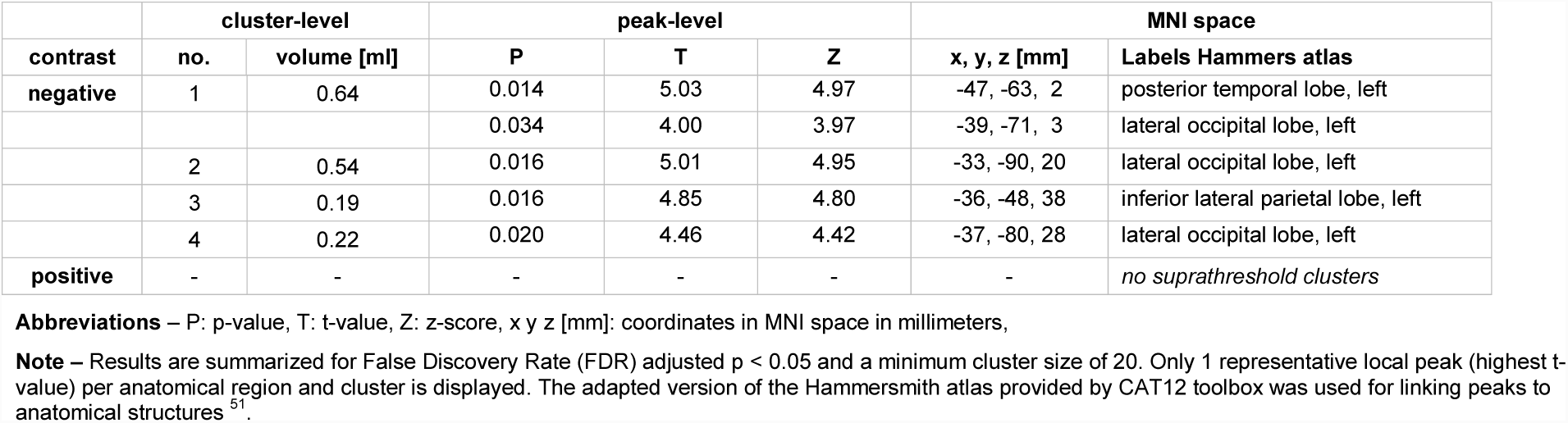
Significant clusters resulting from voxel-wise analysis of plasma cortisol against gray matter volume (corrected for age, TIV; n = 556).

**Figure 3.**
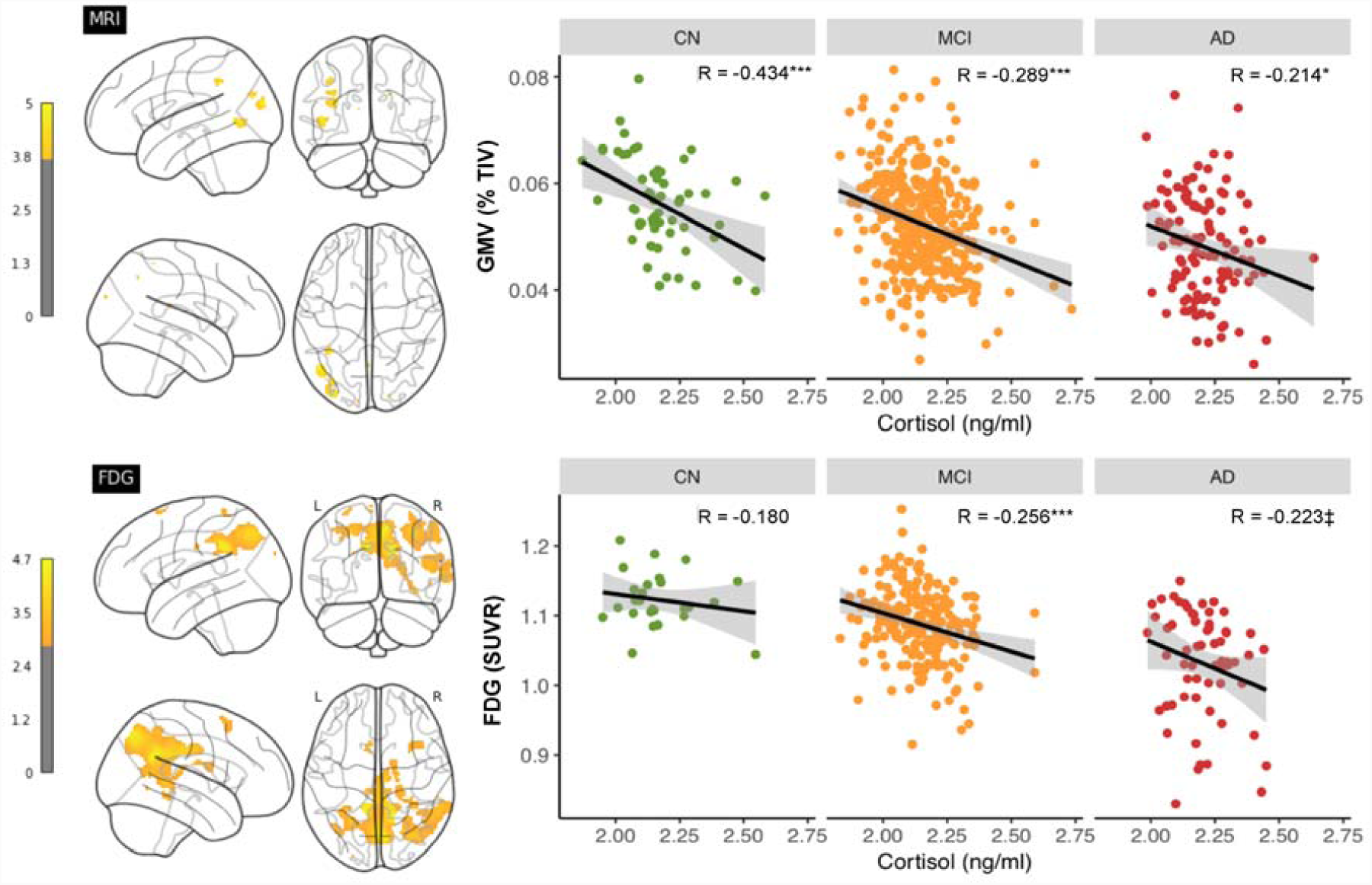
Voxel-wise relationships between plasma cortisol, gray matter volume and glucose metabolism. Top panel shows whole-brain analysis between cortisol and gray matter volume (MRI) and bottom panel between cortisol and glucose metabolism (FGD). Statistical maps (left side) are depicted on a glass brain. Color scales represents t-values starting at p < 0.05, corrected using false discovery rate (FDR). Scatter plots (right side) depict relationships between plasma cortisol and total GMV (measured as % total intracranial volume [TIV]) and average FDG relative standardized uptake value (SUVr) within the corresponding cortisol meta ROI. Dots represent individual data points, with cognitively normal [CN] in green, mild cognitive impairment [MCI] in orange and Alzheimer disease [AD] patients in red. Lines indicate linear trends within each diagnostic group and gray area the standard deviation. Correlation coefficients are provided in the top right for each diagnostic group. Significance levels: ***p ≤ 0.001, **p ≤ 0.01, * p ≤ 0.05, ^‡^ p ≤ 0.1.

Stratification by diagnostic group indicated that the negative relationship between plasma cortisol and gray matter volume in the respective cortisol meta ROI was significant in each diagnostic group (Figure 3 and Table 2B, correlation analysis). There was no significant interaction between diagnostic group and plasma cortisol (*F* [2,550] = 0.292, *p* = 0.747, n = 556).

After adjusting for covariates (age and gender), results were maintained with regard to the entire sample and diagnostic groups (Table 2B, partial correlation analysis).

#### 3.3.2 FDG

At the voxel level, higher plasma cortisol levels were related to lower FDG SUVr mainly within lateral temporo-parietal regions, posterior cingulate, and medial parietal regions including the precuneus of both hemispheres across the entire sample (Figure 3 and Table 4). There were no significant positive relationships between plasma cortisol levels and glucose metabolism at the given threshold.

**Table 4.** Significant clusters resulting from voxel-wise analysis of plasma cortisol against FDG (corrected for age; n = 288).

| contrast | cluster-level |  | peak-level |  |  | MNI space |  |
| --- | --- | --- | --- | --- | --- | --- | --- |
|  | no. | volume [ml] | P | T | Z | x y z [mm] | Labels Hammers atlas |
| negative | 1 | 25.6 | 0.017 | 4.72 | 4.63 | 14, -54, 28 | superior parietal gyrus, right |
|  |  |  | 0.017 | 4.44 | 4.36 | -12, -44, 34 | superior parietal gyrus, left |
|  |  |  | 0.018 | 3.98 | 3.93 | 8, -36, 30 | posterior cinguli gyrus, right |
|  |  |  | 0.028 | 3.58 | 3.54 | -4, -36, 40 | posterior cinguli gyrus, left |
|  |  |  | 0.034 | 3.36 | 3.33 | -33, -50, 42 | inferior lateral parietal lobe, left |
|  | 2 | 3.54 | 0.025 | 3.70 | 3.65 | 58, -42, 2 | posterior temporal lobe, right |
|  |  |  | 0.038 | 3.18 | 3.15 | 60, -52, 22 | inferior lateral parietal lobe, right |
|  | 3 | 7.43 | 0.026 | 3.69 | 3.65 | 34, -72, 48 | inferior lateral parietal lobe, right |
|  |  |  | 0.038 | 3.21 | 3.18 | 36, -74, 36 | lateral occipital lobe, right |
|  | 4 | 0.25 | 0.032 | 3.44 | 3.41 | 34, -40, -8 | posterior temporal lobe, right |
|  | 5 | 0.79 | 0.036 | 3.28 | 3.25 | 24, -30, 14 | insula, right |
|  |  |  | 0.038 | 3.22 | 3.19 | 30, -38, -4 | posterior temporal lobe, right |
|  | 6 | 0.70 | 0.039 | 3.16 | 3.13 | 17, -10, 23 | caudate nucleus, right |
|  | 7 | 0.74 | 0.039 | 3.14 | 3.11 | 32, 16, 60 | middle frontal gyrus, right |
| positive | - | - | - | - | - | - | *no suprathreshold clusters* |

**Abbreviations** – FDG: 2-[F-18]-fluoro-2-deoxy-D-glucose, SUVr: relative standard uptake value (reference region: brain parenchyma), P: p-value, T: t-value, Z: z-score, x y z [mm]: coordinates MNI space in millimeters
**Note** – Results are summarized for False Discovery Rate (FDR) adjusted $p < 0.05$ and a minimum cluster size of 20. Only 1 representative local peak (highest t-score) per anatomical region and cluster is displayed. The adapted version of the Hammersmith atlas provided by CAT12 toolbox was used for linking peaks MNI to anatomically labeled structures (Hammers et al., 2003).

Follow-up stratification by diagnostic group indicated that the negative relationship between plasma cortisol and glucose metabolism in the respective cortisol meta ROI was only significant in the MCI group (Figure 3 and Table 2B, correlation analysis). There was, however, no significant interaction between diagnostic group and plasma cortisol (*F* [2,282] = 0.519, *p* = 0.596, n = 288).

Adjustments for covariates (age and APOE4) did not essentially change the results in the entire sample and within diagnostic groups (Table 2B, partial correlation analysis).

## 4 Discussion

Findings of this study show detrimental effects of plasma cortisol on cerebral glucose metabolism and gray matter volume across the AD spectrum, with a somewhat divergent distribution. Increased plasma cortisol was related to lower glucose metabolism, mainly in medial parietal and lateral temporo-parietal regions. In addition, plasma cortisol was inversely associated with gray matter volume, mainly in lateral temporo-parieto-occipital regions and in the hippocampus. Our results are consistent with the notion that HPA axis activation could exacerbate pathological processes associated with AD and increase the vulnerability to AD development.

### 4.1 Relationship between plasma cortisol and cerebral glucose metabolism

We found that higher overnight fasting plasma cortisol was associated with lower cerebral glucose metabolism across the AD spectrum. This adverse effect was mainly seen within lateral temporo-parietal, medial parietal, and posterior cingulate brain regions. The finding converges with data from animal models, indicating that glucocorticoids may inhibit cerebral glucose metabolism in the hippocampus and other brain regions ^18, 37^. Using FDG PET, previous studies on Cushing’s syndrome patients have further related increased cortisol levels to global and regional patterns of perturbed cerebral metabolism ^19, 20^. Conversely, De Leon and colleagues ^38^ demonstrated that glucose metabolism was specifically reduced in the hippocampus in normal older individuals in response to hydrocortisone administration. Our findings things corroborate the notion that regional glucose metabolism is altered as a function of circulating plasma cortisol.

The inverse relationship between plasma cortisol and glucose metabolism was found in AD signature regions in prodromal and clinical disease stages. Physiological mechanisms underlying the observed effect remain unclear. There is, however, evidence that higher levels of glucocorticoids could haste existing AD pathogenesis ^1, 39^. Elevated glucocorticoid levels were previously linked to increased Aβ accumulation and tau hyperphosphorylation in rodents ^40, 41^ as well as global Aβ burden in humans ^42^. The typical hypometabolic pattern, similarly exhibited by MCI and AD patients ^15, 16, 43^, is suggested to evolve from synergistic interactions of Aβ- and tau-related pathways ^44^. Given these findings it appears that metabolic dysfunction could be aggravated by the impact of stress hormones on existing AD pathologies.

Interestingly, a moderate positive relationship between plasma cortisol levels and FDG uptake was found in the small group of cognitively normal individuals (n = 26), selectively within the pre-defined FDG AD signature ^32^. The unexpected result could potentially reflect neural compensation or aberrant neural activation in the face of neurotoxic insults. For example, a previous study reported higher basal cerebral metabolism as a function of neuropathological burden in patients with MCI ^45^. While compensatory brain responses in AD vulnerable regions could serve to protect higher order brain function, pathological neural overexcitation may instead confer an increased risk of clinical progression ^46^. Further evaluation of this finding and associated mechanisms in larger longitudinal samples is warranted.

### 4.2 Relationship between plasma cortisol and gray matte volume

Higher plasma cortisol levels were associated with lower gray matter volume in the hippocampus and other cortical areas. At the given statistical threshold, the effect was mainly found in restricted left lateral temporal-parietal-occipital regions, with a spatial pattern somewhat distinct from the one found for cerebral metabolism. The finding parallels observations, suggesting that circulating glucocorticoids are associated with volume reduction in widespread brain regions, often comprising but not limited to the hippocampus ^8-10, 47^. The adverse relationship between plasma cortisol and regional volume was found even in cognitively normal individuals, similar to previous reports ^11,47^. One possible explanation might be that circulating glucocorticoids may affect brain structure via several damaging pathways, for example, through synapse remodeling ^48, 49^. Such mechanisms work prior to and independent of AD pathology and may increase the vulnerability of the brain to aging and disease.

### 4.3 Synopsis and outlook

Taken together, our findings are convergent with the idea that HPA axis activation may be a vigorous mechanism that accelerates AD pathogenesis. The study shows detrimental relationships between stress hormone levels and brain integrity within regions sensitive to AD pathology across the AD spectrum. Moreover, we replicate the presence of higher cortical levels in AD patients compared to healthy older adults ^50^. A similar effect was not detectable in MCI patients, albeit reported previously using cortisol measures in CSF ^7^. Clinical or lifestyle interventions could help to reduce elevated stress hormones levels at early disease stages and have a positive impact on disease progression, a hypothesis to be evaluated in future studies.

### 4.4 Strengths and limitations

Our study has strengths and limitations. The present findings are based on the large and well-characterized ADNI cohort with standardized operation procedures that are publicly available. As a result, we were able to investigate plasma cortisol levels across the spectrum of AD, from cognitively normal to mildly and clinically impaired older individuals. However, the use of ADNI is also a limitation, as our findings are not independent from other observation in the same cohort (Toledo et al 2013). Furthermore, the cross-sectional nature of assessments precludes conclusions on the temporal sequence between HPA axis activation and brain injury associated with AD pathogenesis. Longitudinal studies including biomarkers of neuropathology will be needed to further evaluate cortisol-associated effects. Also, single assessments of cortisol levels might not only reflect longer-term effects of high cortisol exposure, as opposed to more dynamic measures over an extended time period. It therefore remains an open question if our findings are mediated by the long-term effects of circulating cortisol or by other factors that influence daily peaks in cortisol. Finally, our findings are likely weighted towards prodromal disease stages, with MCI patients being the largest participant groups. Forthcoming studies could assess more fine-grained effects of plasma cortisol exposure across the complete AD continuum.

## 5 Conclusion

The present study demonstrated that higher plasma cortisol levels is related to reduced glucose metabolism in AD signature regions across the AD spectrum. Higher cortisol is also related to reduced regional gray matter volume, with partially overlapping spatial regions across the brain. The detrimental relationship between plasma cortisol and glucose metabolism were most pronounced in prodromal and clinical stages of AD. The findings suggest that HPA axis activation could aggravate existing AD pathological processes and thereby accelerate AD progression.

## 6 Acknowledgments

We sincerely thank the ADNI research group for their contributions to this work and Dr. Etienne Vachon-Presseau (Northwestern University, Chicago) and Gloria Benson (Charité – Universitätsmedizin, Berlin) for valuable comments on the manuscript.

## 6.1 Study funding

The present study was supported by the Hans Gerhard Creutzfeldt scholarship (FKZ CSB II, 01EO1301 TP T2) and the NeuroCure Female Postdoctoral research fellowship (Exc 257/2). Data collection and sharing for this project was funded by the Alzheimer’s Disease Neuroimaging Initiative (ADNI) (National Institutes of Health Grant U01 AG024904) and DOD ADNI (Department of Defense award number W81XWH-12-2-0012). ADNI is funded by the National Institute on Aging, the National Institute of Biomedical Imaging and Bioengineering, and through generous contributions from the following: AbbVie, Alzheimer’s Association; Alzheimer’s Drug Discovery Foundation; Araclon Biotech; BioClinica, Inc.; Biogen; Bristol-Myers Squibb Company; CereSpir, Inc.; Eisai Inc.; Elan Pharmaceuticals, Inc.; Eli Lilly and Company; EuroImmun; F. Hoffmann-La Roche Ltd and its affiliated company Genentech, Inc.; Fujirebio; GE Healthcare; IXICO Ltd.; Janssen Alzheimer Immunotherapy Research & Development, LLC.; Johnson & Johnson Pharmaceutical Research & Development LLC.; Lumosity; Lundbeck; Merck & Co., Inc.; Meso Scale Diagnostics, LLC.; NeuroRx Research; Neurotrack Technologies; Novartis Pharmaceuticals Corporation; Pfizer Inc.; Piramal Imaging; Servier; Takeda Pharmaceutical Company; and Transition Therapeutics. The Canadian Institutes of Health Research is providing funds to support ADNI clinical sites in Canada. Private sector contributions are facilitated by the Foundation for the National Institutes of Health (www.fnih.org). The grantee organization is the Northern California Institute for Research and Education, and the study is coordinated by the Alzheimer’s Disease Cooperative Study at the University of California, San Diego. ADNI data are disseminated by the Laboratory for Neuro Imaging at the University of Southern California.

## 6.2 Author contributions

Miranka Wirth: study concept and design, data analysis, interpretation of study results, and manuscript drafting

Catharina Lange: data analysis, interpretation of study results, manuscript drafting, final manuscript review

Willem Huijbers: study concept and design, data analysis, interpretation of study results, manuscript drafting, final manuscript review

## 6.3 Disclosures

Miranka Wirth: nothing to disclose.

Catharina Lange: nothing to disclose.

Willem Huijbers: nothing to disclose.

